# Slow kinesin-dependent microtubular transport facilitates ribbon synapse assembly in developing cochlear inner hair cells

**DOI:** 10.1101/2024.04.12.589153

**Authors:** Roos Anouk Voorn, Michael Sternbach, Amandine Jarysta, Vladan Rankovic, Basile Tarchini, Fred Wolf, Christian Vogl

**Author notes:** UCB Pharmaceuticals, 1070 Brussels, Belgium.

## Abstract

Sensory synapses are characterized by electron-dense presynaptic specializations, so-called synaptic ribbons. In cochlear inner hair cells (IHCs), ribbons play an essential role as core active zone (AZ) organizers, where they tether synaptic vesicles, cluster calcium channels and facilitate the temporally-precise release of primed vesicles. While a multitude of studies aimed to elucidate the molecular composition and function of IHC ribbon synapses, the developmental formation of these signalling complexes remains largely elusive to date. To address this shortcoming, we performed long-term live-cell imaging of fluorescently-labelled ribbon precursors in young postnatal IHCs to track ribbon precursor motion. We show that ribbon precursors utilize the apico-basal microtubular (MT) cytoskeleton for targeted trafficking to the presynapse, in a process reminiscent of slow axonal transport in neurons. During translocation, precursor volume regulation is achieved by highly dynamic structural plasticity – characterized by regularly-occurring fusion and fission events. Pharmacological MT destabilization negatively impacted on precursor translocation and attenuated structural plasticity, whereas genetic disruption of the anterograde molecular motor Kif1a impaired ribbon volume accumulation during developmental maturation. Combined, our data thus indicate an essential role of the MT cytoskeleton and Kif1a in adequate ribbon synapse formation and structural maintenance.

## Introduction

Sensory perception requires a sophisticated encoding system that faithfully conveys sudden changes in environmental conditions with utmost temporal precision. To facilitate this challenging task, sensory receptor cells in the vertebrate eye and ear are equipped with presynaptic specializations – ‘synaptic ribbons’ – which tether glutamate-filled synaptic vesicles (SVs) and cluster presynaptic Ca^2+^ channels at the releases site (Moser et al., 2019). Besides their structural role as the main active zone (AZ) scaffold, sensory ribbons are thought to facilitate SV priming and act as ‘conveyor belts’ that mediate vesicular replenishment during periods of sustained activity (Joselevitch and Zenisek, 2020; LoGiudice et al., 2008; Vaithianathan et al., 2016). While extensive efforts have been made to dissect the molecular composition and function of these high-throughput synapses, their developmental assembly remains largely elusive to date.

In the cochlea, ribbon synapse formation involves the accumulation of multiple small ribbon precursor spheres at the afferent contacts of late embryonic inner hair cells (IHCs) (Michanski et al., 2019; Sobkowicz et al., 1986). These precursors are most likely generated via cytosolic RIBEYE auto-aggregation (Magupalli et al., 2008; Schmitz et al., 2000), which occurs within the basolateral IHC compartment. Such ‘free-floating’ ribbon precursors can occur at significant distances to the AZ and have previously been observed not only in auditory IHCs, but also pinealocytes and retinal photoreceptors (Hermes et al., 1992; Regus-Leidig et al., 2009; Spiwoks-Becker et al., 2008, 2004). Across all systems, floating precursors were shown to tether a cohort of SVs and consist not only of the main scaffold RIBEYE, but also contain other AZ proteins such as Piccolino, a short splice variant of Piccolo (Michanski et al., 2023, 2019; Regus-Leidig et al., 2013). Hence, these precursors can be considered as presynaptic ‘building blocks’ for rapid AZ establishment or supplementation at nascent afferent contacts. While their mode of transport towards the developing AZ still remains to be established, our previous work revealed close spatial proximity of ribbon precursors and the microtubule (MT) network in murine IHCs and found the MT plus end (+end) -directed molecular motor Kif1a to colocalize with cytosolic ribbon precursors (Michanski et al., 2019). These findings strongly suggest MT-based precursor translocation during IHC development; yet, all work to date has been performed in fixed samples, and therefore dynamic precursor movement or targeted transport remains to be unambiguously demonstrated.

In the present study, we analyze ribbon precursor trafficking in the murine organ of Corti using a comprehensive live-cell and fixed-tissue imaging approach. We first establish MT network polarization in fixed tissue, to then visualize precursor movement along MTs and perform real-time tracking analyses on fluorescently-labelled ribbon precursors of organotypically-cultured IHCs *in vitro*. Additionally, we employ pharmacological manipulation to probe the role of the MT cytoskeleton in precursor translocation and investigated the function of Kif1a in precursor trafficking by analyzing auditory brainstem responses (ABRs), synapse counts and ribbon volumes in IHCs of *Kif1a leg dragger* (*Kif1a^lgdg^*) mouse mutants, which harbor a L181F point mutation in the Kif1a motor domain that leads to functional impairment. In line with our hypothesis, we find that acute pharmacological destabilization of the IHC cytoskeleton alters ribbon precursor dynamics, volume acquisition and structural plasticity, whereas genetic disruption of Kif1a function leads to attenuated ABRs and reduced ribbon volumes in *Kif1a^lgdg^* mutants.

Together with our companion paper (Hussain et al.), which assesses ribbon precursor trafficking in developing zebrafish lateral-line neuromast hair cells, our data provide the first direct experimental evidence of targeted MT-based and Kif1a-dependent ribbon precursor trafficking in synaptogenesis and maturational refinement of IHC ribbon synapses.

## Results

### Developing IHCs exhibit a highly polarized and strongly acetylated apico-basal MT cytoskeleton

MTs are delicate and highly dynamic cytoskeletal elements that provide structural stability as well as cell shape and additionally mediate targeted intracellular transport. Consisted with other polarized cell types and in line with previous work (Akhmanova and Kapitein, 2022; Steyger et al., 1989), we find that in IHCs, the MT network was oriented along the apico-basal axis of the cell, exceedingly dense and highly branched (Figure 1A). Moreover, large parts of the MT cytoskeleton were found to be acetylated – with an apical-to-basal gradient (Figure 1B). In support of a more stabilized MT network at the IHC apex, immunolabelling of the non-centrosomal MT minus-(–)end binding protein CAMSAP2 (Tanaka et al., 2012) confirmed that the vast majority of the MT strands are indeed polarized from the apical cell pole towards the basolateral compartment, as CAMSAP2 immunofluorescence was largely restricted to the IHC neck region (Figure 1C-C”).

**Figure 1:**
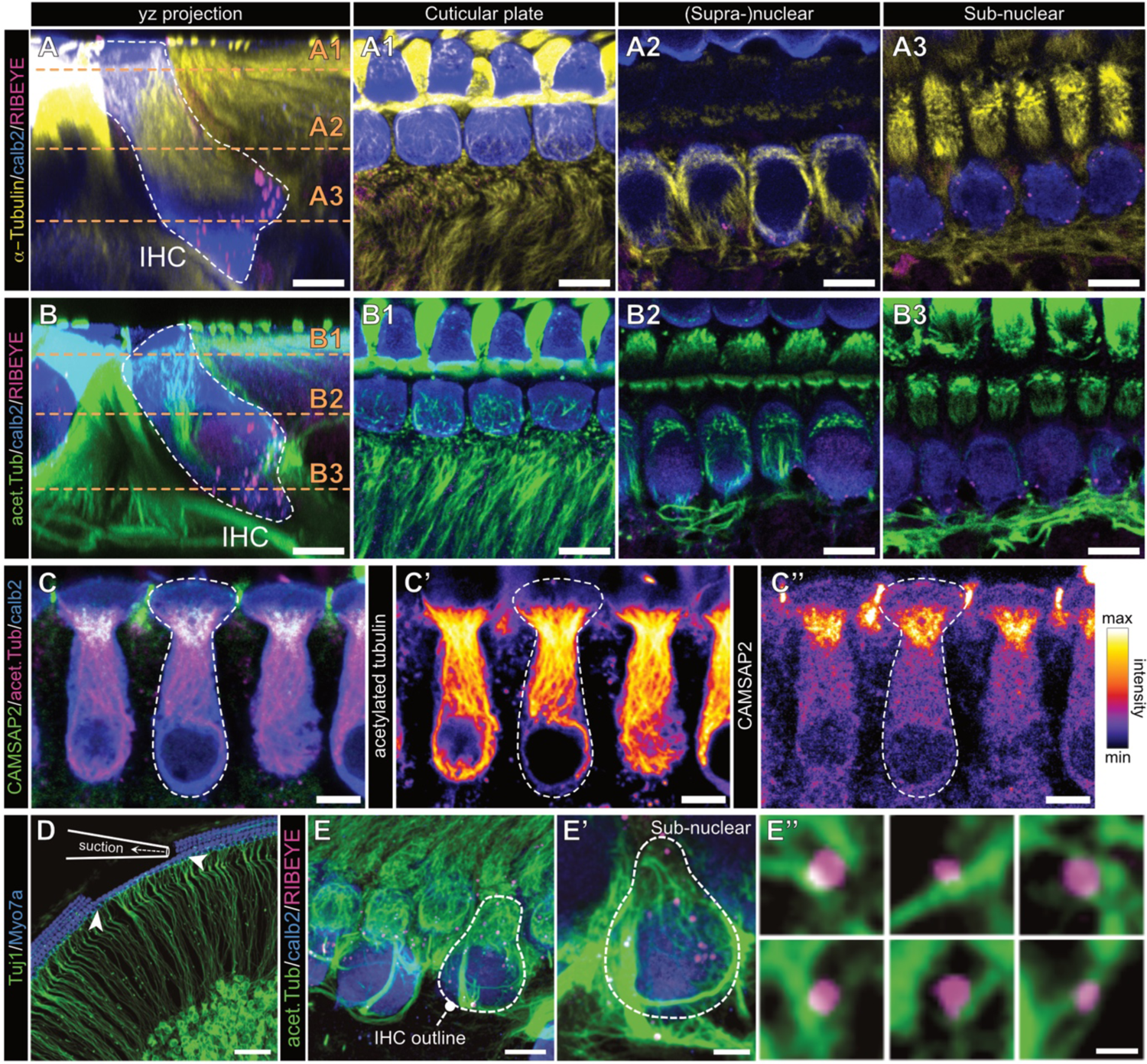
The apico-basally polarized IHC microtubule cytoskeleton is highly acetylated. **A** Representative confocal yz-projection of an immunohistochemical staining of the MT cytoskeleton in acutely dissected organs of Corti of early postnatal mice (P7). Indicated is the IHC outline (white dashed line), labeled for α-tubulin (yellow), ribbons (RIBEYE, magenta) and IHC context (Calretinin, blue). **A1-A3** different axial sections of IHC and MT labeling from A, displaying the MT cytoskeleton localization within the IHCs and in the surrounding tissue. **B** Representative confocal images and sectioning (**B1-3**) as in **A**, but for immunolabeling of acetylated tubulin (green). Please note that in A1-3 and B1-3 the intensity levels of the tubulin channels have been adjusted for optimal visibility. **C** Immunohistochemical labeling of MT –end binding protein CAMSAP2 (green), and acetylated MT strands (magenta) within IHCs (Calretinin, blue), in acutely dissected organs of Corti (P12). **C’** Acetylated tubulin strands reach from the cellular apex into the basolateral synaptic area. **C’’** CAMSAP2 labeling is specifically localized in the apical IHC just below the cuticular plate. **D** Schematic depiction of the mechanical cleaning technique used to remove OHCs, inner pillar cells and phalangeal cells to facilitate unobstructed access to the row of IHCs. Hair cells are labelled for Myo7a (blue), spiral ganglion neurons for β3-tubulin/Tuj1 (green). **E-E’** Immunohistochemical labeling of mechanically-cleaned organotypic cultures of the organ of Corti, stained for IHCs (Calretinin, blue), acetylated tubulin (green) and ribbon precursors (RIBEYE, magenta). **E’’** Higher magnification single confocal sections of ribbon precursors colocalizing with acetylated MT strands. Scale bars: A-B” & D, 5 μm; C, 50 μm; D’, 2.5 μm; D”, 0.5 μm.

Reproducible MT immunolabelling in the basolateral compartment of developing IHCs proved a surprisingly difficult task. This is likely due to the very high density of tubulin strands in the surrounding supporting cells (Figure 1A,B). This configuration might act as an ‘antibody sink’ that leads to local depletion and ultimately decreased labelling intensity in the comparatively much less tubulin-expressing IHCs. We resolve this issue with a mechanical cleaning approach that is commonly used for patch clamp electrophysiology (Figure 1D). Here, the three rows of outer hair cells (OHCs) and supporting cell layers were physically removed with a glass micropipette prior to tissue fixation and immunostaining. This approach facilitated antibody accessibility to the otherwise deeply-embedded IHC basolateral poles and allowed for improved visualization of ribbon precursor-MT interactions. In line with our previous work (Michanski et al., 2019), we found numerous occasions of direct appositions between ribbon precursors and tubulin strands reaching deep into the presynaptic compartment that are compatible with active precursor transport along apico-basally polarized tubulin strands (Figure 1E-D”). Moreover, given the directionality of MT growth, a MT +end directed molecular motor – such as the previously implicated kinesin-3 Kif1a (Michanski et al., 2019) – can be suspected to facilitate this process.

### Ribbon precursors translocate along MTs in living IHCs

To test if ribbon precursor translocation to the presynapse indeed requires active MT-based transport, we devised a novel live-cell approach for *in situ* imaging of this process in mammalian IHCs (Figure 2; Supplemental Figure 2-1): Similar to the above-described MT immunolabelling approach, we used mechanical cleaning to optimize the strength and signal-to-noise ratio of the MT labeling via the fluorogenic live-cell dye SPY555-tubulin. Following method optimization, we conducted long-term live-cell imaging experiments on organotypic cultures prepared from mice that were virally-injected with a RIBEYE-GFP encoding AAV one day prior. This approach faithfully labelled ribbon precursors and enabled live-cell tracking of individual precursor movement alongside filamentous MT strands in living IHCs (Figure 2A,B).

**Figure 2.**
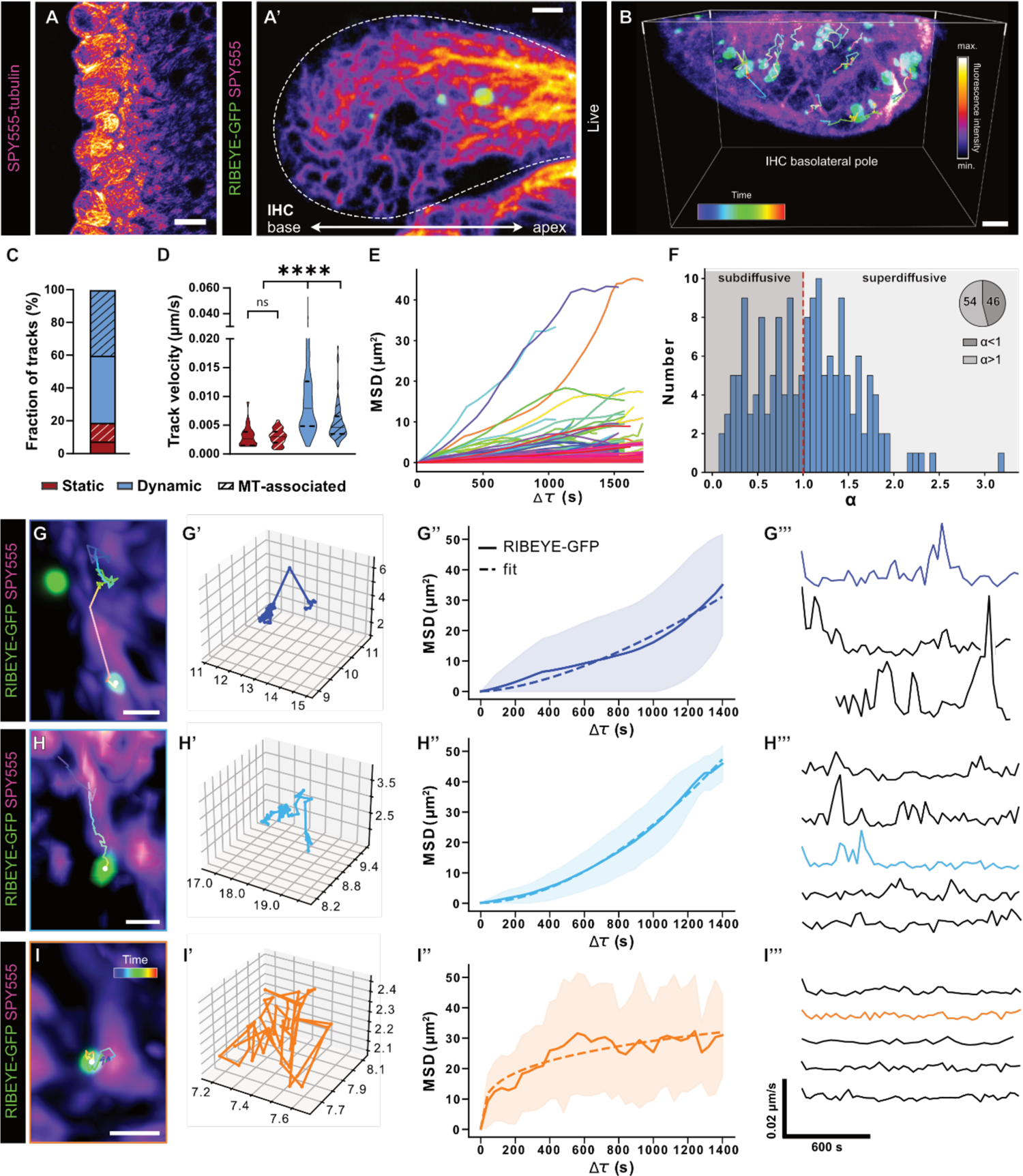
Visualizing MT-based ribbon precursor transport in living IHCs. **A** Representative confocal live-cell image of the IHC row of an organotypic culture labeled with SPY555-tubulin, in which the outer hair cells, inner pillar cells and phalangeal cells have been mechanically removed. **A’** Higher magnification live-cell image of an exposed IHC, labeled with SPY555-tubulin with surrounding tissue cleaned, expressing RIBEYE-GFP (green). **B** Three-dimensional reconstruction of live-cell timelapse imaging of the basolateral compartment of a RIBEYE-GFP transduced IHC additionally labelled with SPY555. Single particle tracking of ribbon precursors within the basolateral IHC reveals highly dynamic displacements. Trajectories are color-coded for time. Total imaging time: 40 min. **C** The majority of traced ribbon precursors were classified as mobile (displacing >1 µm in 30 min). Half of the mobile population could be detected to displace along MTs. **D** Although static ribbon precursors showed low displacement over time, precursors did undergo moderate spatial fluctuation, leading to a low average track velocity. While both mobile populations showed a considerably higher average velocity than the static precursors, remarkably, the track velocity of precursor displacement independent from MTs was significantly higher than of MT-associated precursors. **E** Combined plot of the mean squared displacement (MSD) of all MT-associated ribbon precursor trajectories, indicative of multiple types of motion. **F** Distribution of the exponent α, extracted from the MT-associated precursor tracks, where α=1 equals a diffusive or Brownian motion, α<1 indicates subdiffusion for confined motion, and α >1 directed transport. **G, H, I** Example trajectories of ribbon precursors in association with the MT cytoskeleton. Three main types of motion could be observed: (**G**) stop-and-go displacement, including rapid long-distance traversing jumps, as well as intermittent periods of near static behavior, (**H**) slow continuous, near linear progressive motion in a targeted fashion along the MT strand and (**I**) confined motion in place but attached to the MT network. Of the three main MT-associated motion types, a three-dimensional representation is plotted (**G’, H’, I’**), as well as the MSD of the respective trajectories (**G’’, H’’, I’’**) – please note that individual scales have been adapted for optimal visibility of the respective trajectory. During precursor displacement, we detected significant velocity fluctuations of which representative sample traces are shown per motion subtype (**G’’’, H’’’, I’’’**). Illustrated examples indicated by consistent coloring and color-coded for time. Values represented as violin plots, with medians and the 25% and 75% interquartile range indicated with solid and dashed lines respectively. Statistical significance: Kruskal-Wallis. ****p<0.0001. N=5, n=8. Scale bars: A-B, 10 μm; A’, B’-C, 2 μm; G-I 1 μm.

Our live-cell imaging experiments indicated that ∼20% of ribbon precursors remained in a stable position during the 40 min observation period, whereas the remaining ∼80% exhibited various degrees of mobility. Interestingly, roughly half of this mobile fraction appeared to translocate along SPY555-labelled MTs (Figure 2C). Velocity analysis of the different mobile precursor populations revealed that non-MT-associated ribbons displaced at average velocities of 0.0995 ± 0.0006 μm/s, while MT-associated precursors moved with significantly *lower* velocities (0.0055 ± 0.0003 μm/s; P<0.0001; Figure 2D). To test if the observed motion is directional, we employed mean squared displacement (MSD) analysis of all MT-associated ribbons. Here, the extracted exponent α can be used to distinguish diffusive/non-directional behavior (α=1) from subdiffusive/confined (α<1) and superdiffusive/targeted (α>1) motion. This analysis revealed that a significant fraction of ribbon precursors (∼54%) underwent targeted transport (Figure 2E-F). In fact, upon closer inspection of our live-cell data, we found three distinct types of MT-bound motility: (i) a clearly directional ‘saltatory’ mode that was characterized by intermittent periods of rapid movements in-between extended periods of confinement – a behavior indicative of interrupted motor-based transport events (Figure 2G-G’’), (ii) a gradual/continuous mode that also appeared highly directional (Figure 2H-H’’) and (iii) a seemingly non-directional confined mode (Figure 2I-I’’). Importantly, precursors of both directionally-displacing categories presented with supralinear MSDs, indicative of targeted transport along the associated MT tracks. Remarkably, the gradual/continuous mode appeared to dominate as the preferred mode of precursor translocation (∼74%).

Since the observed velocities of MT-associated precursors resided in the range of slow axonal transport of cytosolic protein aggregates in neurons (Lasek et al., 1984) and the frequency and duration of MT-association has been suggested to ultimately determine transport velocity, we assessed the velocity profiles of individual MT-associated precursors and found that both, the saltatory and continuous progressing trajectories exhibited a high degree of velocity fluctuation that was characterized by the regular occurrence of defined peaks of increased speed (Figure 2G’’’-I’’’). Thus, these data are in line with a ‘stop-and-go’ processivity that generates anterograde movement.

### Ribbon precursor volume is dynamically modified through bi-directional plasticity events

In addition to a basic characterization of precursor mobility, our live-cell imaging experiments allowed the detailed analysis of ribbon precursor structural plasticity during the observation period (Figure 3). Here, lineage tracing analysis revealed complex patterns of *bidirectional* plasticity events that dynamically modified precursor volume during their journey through the IHC cytoplasm (Figure 3A). Specifically, we observed frequent occurrences of precursor fusions (Figure 3B-B’) and – to our surprise – fission events (Figure 3C-C’). Interestingly, the level of interactivity was highly variable between precursors, where some remained structurally stable, whilst others were subject to multiple plasticity events during the total imaging time.

**Figure 3:**
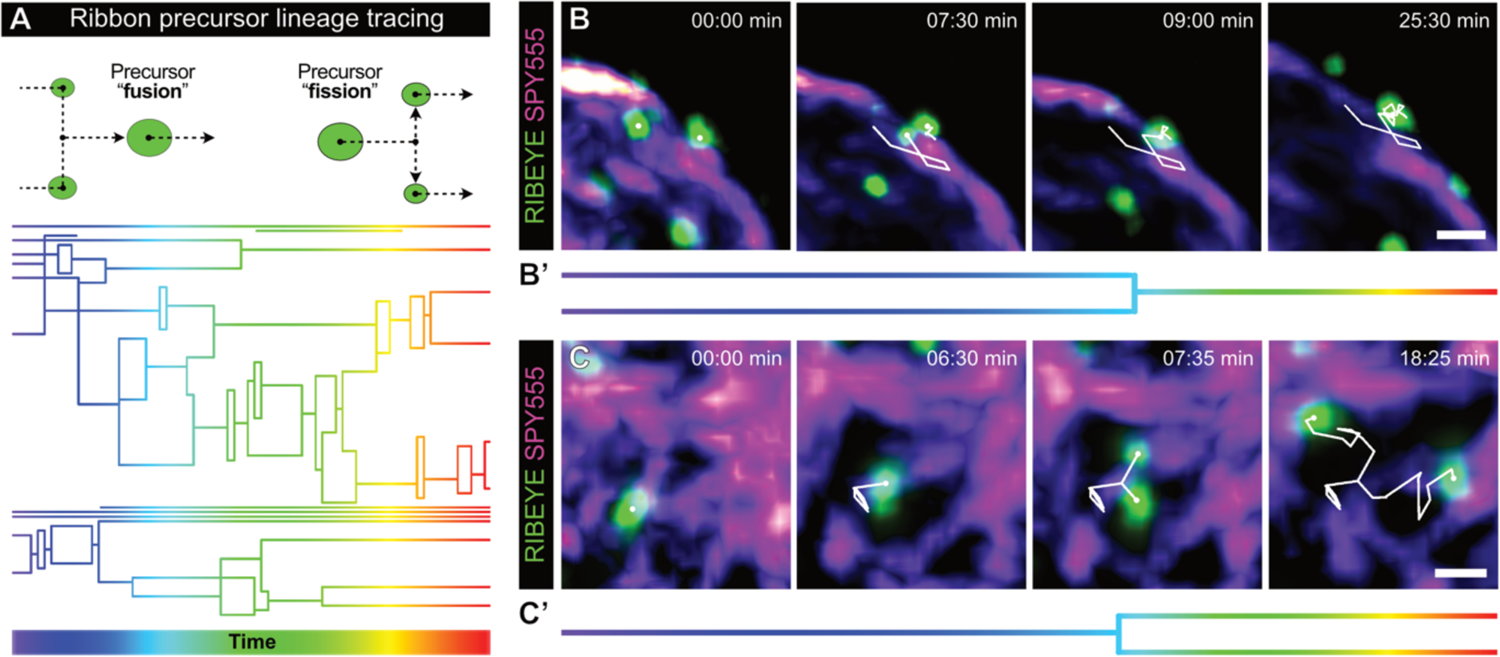
Ribbon precursor volume is dynamically modified by bi-directional structural plasticity. **A** Upper panel: Schematic representation of detected plasticity events using the lineage tracing algorythm (IMARIS). Ribbon precursors could be observed to undergo fusions and fissions. Lower panel: representative example of a lineage tracing graph for all precursors of one IHC, color-coded for time. Total live-cell imaging time: 40 min. As indicated in the upper panel, each horizontal line represents one ribbon precursor at that specific time point. Vertical connections reflect plasticty events. **B,C** Exemplary confocal live-cell imaging frames, displaying different steps in the plastic process of precursors fusion (**B**), and fission of one precursor into two (**C**). **B’,C’** Representations of the plasticity event examples in (**B**) and (**C**) in simplified style of the lineage tracing graphic. Precursors labeled by viral transduction of RIBEYE-GFP; additonal labeling of the MT cytoskeleton by application of SPY555-tubulin. Scale bars: 1 μm.

### Acute nocodazole treatment attenuates ribbon precursor mobility in living IHCs

Our imaging dataset strongly indicates that a significant subset of ribbon precursors indeed utilizes MT tracks for intracellular translocation. To further confirm the role of MTs in precursor transport, we used pharmacology to acutely destabilize the MT cytoskeleton during our live-cell imaging experiments and assess effect on precursor mobility (Figure 4). Methodologically, it can be challenging to destabilize MT when they are – at least partly – stabilized by the paclitaxel-based SPY555-tubulin dye. Therefore, we opted for a different context marking approach and used viral overexpression of RIBEYE-tdTomato in IHCs of Ai32-Vglut3-Cre knock-in reporter mice (Ai32-VC-KI) (Chakrabarti et al., 2022). Here, IHCs express a YFP-tagged channelrhodopsin-2/H134R (ChR2-YFP) in the plasma membrane. Importantly, YFP and ChR2 have distinct excitation/photoactivation spectra, enabling us to use YFP to visualize IHCs without activating ChR2 during our investigation. Organotypic cultures of these mice were prepared and used for timelapse imaging. We then incubated these cultures either with vehicle control (DMSO) or the MT-depolymerizing agent nocodazole and monitored the movement of all RIBEYE-tdTomato containing ribbon precursors per IHC over a 40 min time period (Figure 4A). In line with our above observations, precursor tracing revealed highly dynamic networks of structural plasticity, several aspects of which were affected by the nocodazole treatment: For example, nocodazole significantly reduced precursor velocity (vehicle_median_ 0.0052 μm/s, IQR 0.0033-0.0084; nocodazole_median_ 0.0050 μm/s, IQR 0.0032-0.0080; P=0.0003; Figure 4B) and thus attenuated the total displacement of precursors within their traveled trajectories (vehicle_median_ 0.085 μm, 0.040-0.180; nocodazole_median_ 0.055 μm, IQR 0.027-0.114; p<0.0001; Figure 4C). Strikingly, lineage tracing analysis revealed that upon nocodazole-based MT destabilization, the complexity of precursor tracks was dramatically reduced – indicative of an essential role of the intact MT cytoskeleton for adequate ribbon synapse assembly. In fact, ribbon precursors underwent significantly fewer fusions (vehicle_mean_ 6.4 ± 1.2 events/h; nocodazole_mean_ 2.9 ± 0.7 events/h; p=0.0343; Figure 4D), and fission events (vehicle_mean_ 7.4 ± 1.0 events/h; nocodazole_mean_ 3.2 ± 0.8 events/h; p<0.0085; Figure 4E). As a result, precursors spent more time in stable non-interactive trajectories (vehicle: 30% vs. nocodazole: 49%) and correspondingly less time in trajectories of high structural plasticity (vehicle: 29%, nocodazole: 16%; Figure 4F). Finally, IHCs of nocodazole-treated cultures exhibited slightly increased ribbon volumes (vehicle_median_ 0.81 μm^3^, IQR 0.39-1.62; nocodazole_median_ 1.1 μm^3^, IQR 0.43-2.12; p<0.0001 Figure 4G-G’) and displayed a tendency towards decreased ribbon numbers, although this trend did not pass our criterion for statistical significance (Figure 4H).

**Figure 4:**
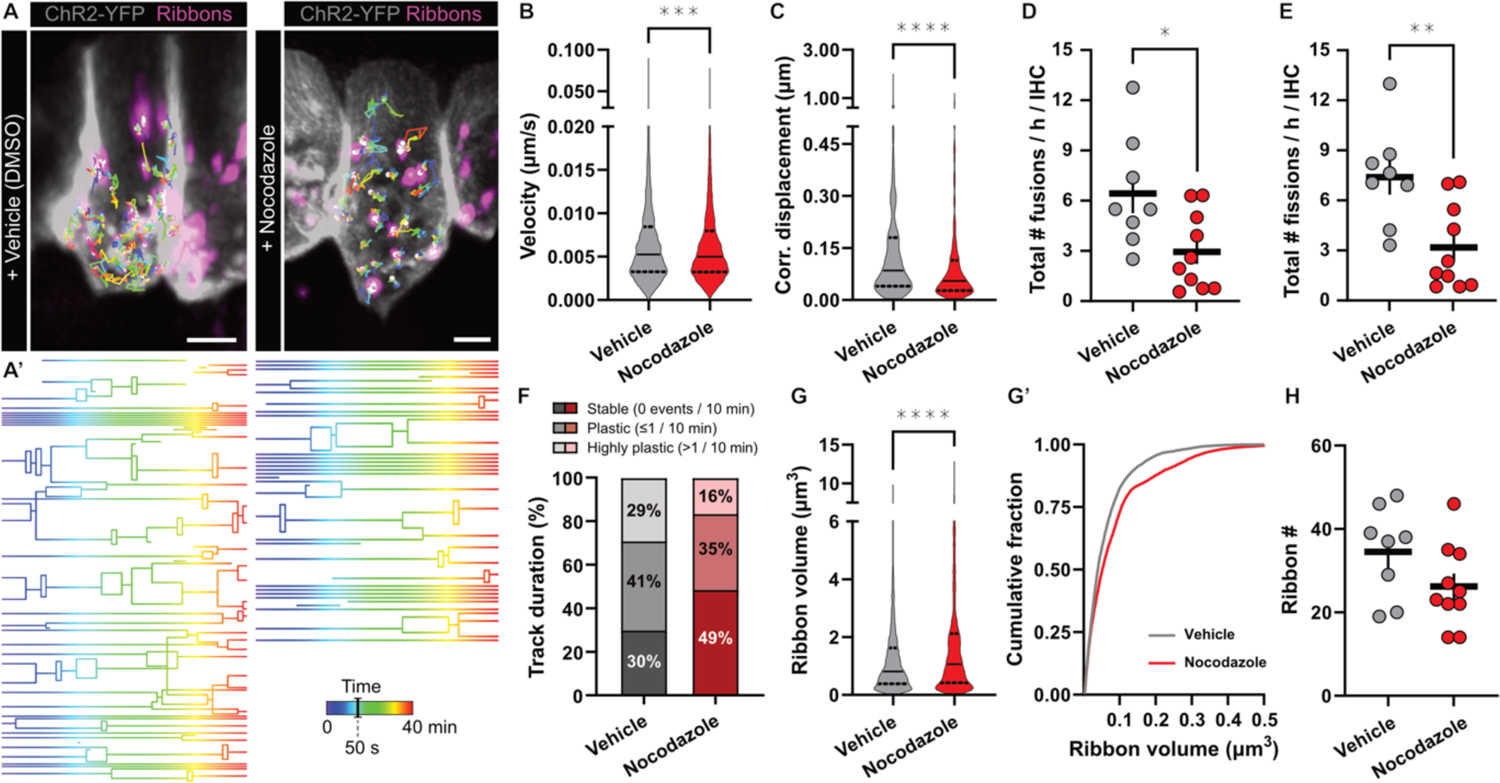
Live-cell IHC ribbon precursor dynamics upon pharmacological disruption of the MT cytoskeleton. **A** Representative live-cell imaging stills of organotypically-cultured IHCs of Ai32-VC-KI mice, with (ChR2-coupled) YFP expression decorating the IHC membrane, and virally-expressed RIBEYE-tdTomato. Ribbon precursor temporal trajectories are color-coded for time. **A’** Graphical representation of the lineage tracing-based ribbon precursor motion over time, illustrating precursor fusion and fission in the cytoplasm. Total imaging time: 40 min. **B** The velocity of individual precursor particles was slightly reduced upon acute treatment with the MT-destabilizing drug nocodazole (1 μM, 3 h). **C** The displacement of ribbon precursors over the course of their trajectories was significantly lower upon nocodazole treatment. Displacement length corrected for the duration of the trajectory, calculated as displacement in 1 minute. **D,E** Interestingly, the frequency of plasticity events within the precursor trajectories was significantly reduced, as precursors were observed to undergo significantly fewer (**D**) fusion as well as (**E**) fission events. **F** The nocodazole-induced reduction in plasticity event frequency resulted in precursors spending an increased percentage of time in individually stable, non-interactive trajectories. On the other hand, the presence of highly dynamic trajectories was reduced. **G-G’** Ribbon precursor volume was increased upon acute nocodazole treatment. **H** The number of ribbon precursors per IHC was not significantly affected by nocodazole treatment. Values represented as violin plots, with medians and the 25% and 75% interquartile range indicated with solid and dashed lines respectively. Statistical significance: Mann-Whitney U. **p<0.01, ****p>0.0001. N=6, n=18. Scale bar: 5 μm.

We next took a closer look at nocodazole effects on precursor mobility and directionality by analyzing MSD and additionally calculating trajectorial asymmetry (Figure 5). Here, purely symmetric particle displacement would indicate random Brownian motion, whereas asymmetric displacement is indicative of targeted transport with a directional bias. In line with our hypothesis of MT-mediated precursor transport, we saw a reduction in precursor MSDs in nocodazole-treated IHCs, yet failed to detect any indications of trajectorial asymmetry (Figure 5A,B). Here, we suspect that the mixed modes of directional and non-directional mobility within the entire ribbon population may compromise our asymmetry analysis and therefore – based on our findings that the main MT-associated precursor fraction undergoes a slow mode of targeted transport – we focused our subsequent analyses on the precursor subpopulation that traveled at speeds below the mean velocity. Indeed, nocodazole application appeared to induce a slight left shift in the velocity distribution (Figure 5C). To investigate this in more detail, we introduced a low (<0.0056 µm/s) and high (>0.01 µm/s) velocity cut-off to differentially analyze slow and fast displacing precursor populations individually. MSD analysis of low velocity precursor trajectories confirmed the suspected loss of directed motion in this population upon nocodazole treatment (Figure 5D). Moreover, the clear trajectorial asymmetry that was found under control conditions – indicative of a biased directionality – was lost upon nocodazole application. These findings hence confirm the notion that targeted precursor transport is facilitated by a slow anterograde process that critically requires an intact MT cytoskeleton. Compatible with this hypothesis, this effect became increasingly less obvious when we expanded our analysis window to include larger fractions of faster-displacing but less directional precursors, thereby ‘diluting’ the slow-directional subpopulation (Supplemental Figure 5-S1). Moreover, when assessing the mobility behavior of high velocity tracks (Figure 5E), we found that, while the MSD was still attenuated by nocodazole treatment, trajectorial asymmetry was absent in both experimental conditions. This behavior is compatible with a combination of (i) fast active transport in multiple opposing directions and/or (ii) a larger contribution of MT-independent non-directional mobility (e.g., during free-floating subdiffusive periods) constituting this latter subpopulation.

**Figure 5:**
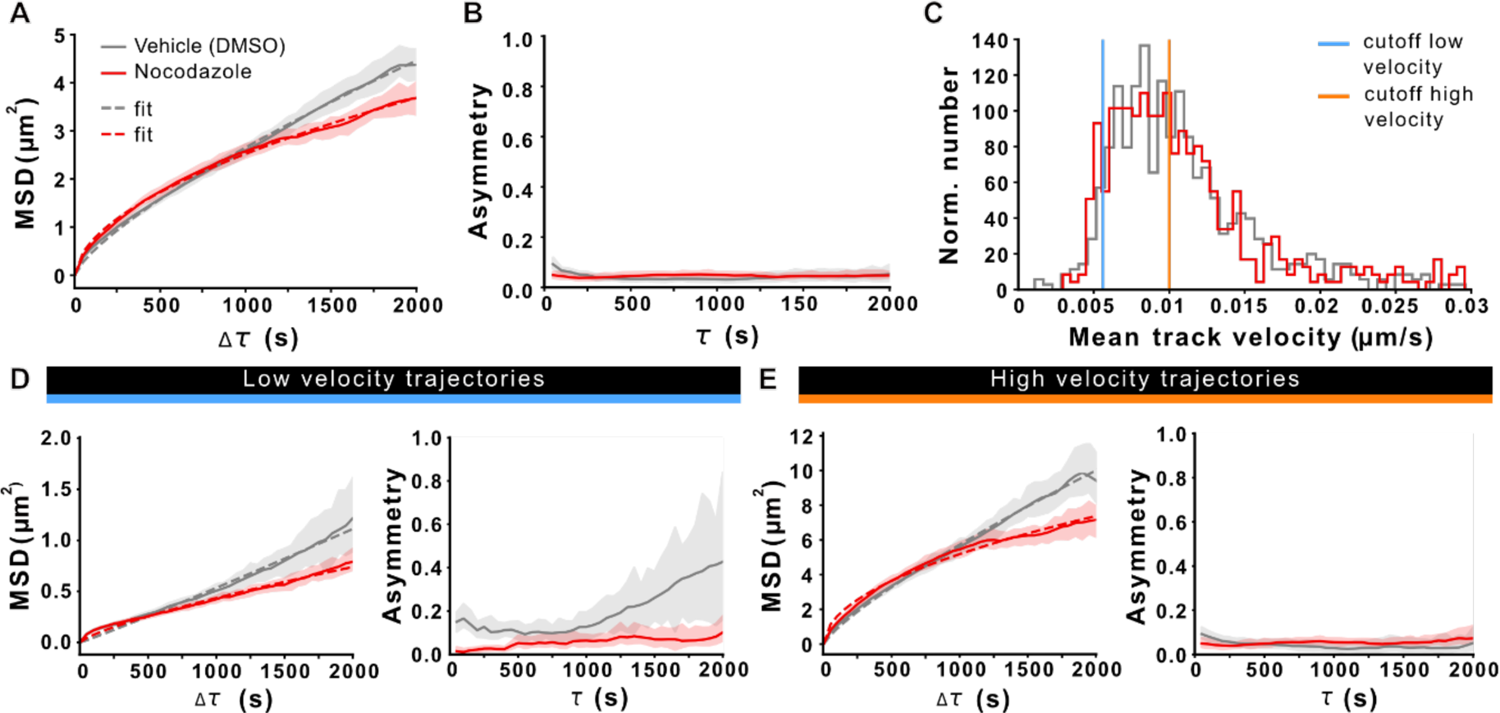
Analysis of three-dimensional ribbon precursor displacement and directionality of motion. **A** Mean square displacement (MSD) of ribbon precursors traced in control conditions (DMSO) and after incubation with nocodazole. N(exp.)=6, n(IHC)=18, n(particles)= 604(DMSO), 462(nocodazole). Nocodazole-induced MT destabilization reduced the MSD. **B** Assessment of the (an)isometry/asymmetry of precursor motion. **C** Distribution of the mean track velocity for precursors in vehicle- and nocodazole-treated IHCs. Indicated are the used cutoffs to selectively analyze trajectories with a low (blue) and high velocity (orange) displacement. **D** MSD analysis of trajectories with a low mean velocity reveals a loss of directed motion upon nocodazole treatment (left panel). Trajectorial asymmetry analysis of slow transport tracks shows a clear directionality for precursors in the vehicle treated condition that is absent in nocodazole-treated IHCs (right panel). n(particles)=40(DMSO), 41(nocodazole). **E** The MSD of high velocity trajectories shows a moderate reduction in directed transport resulting from nocodazole treatment (left panel), but do not show preferential directionality, as apparent from the lack of trajectorial asymmetry (right panel). n(particles)=295(DMSO), 224(nocodazole)

### Genetic disruption of Kif1a impacts hearing and ribbon synapse development

Due to the fact that slow axonal trafficking of soluble synaptic molecules requires short spurts of fast-moving transport in neurons (Tang et al., 2013) and slow bulk transport of choline acetyltransferase in *Drosophila* axons was reported to be a kinesin-dependent process (Sadananda et al., 2012), we revisited our previous hypothesis that Kif1a might be involved in ribbon precursor trafficking (Figure 6). Since *Kif1a-*KO mice die shortly after birth (Yonekawa et al., 1998), we used the viable *Kif1a^lgdg^* mouse model to analyze auditory brain stem responses (ABRs), IHC synapse counts and ribbon volumes. *Kif1a^lgdg^* mice show a progressively deteriorating phenotype that ultimately leads to hindlimb paralysis within 3 - 4 weeks of birth. Therefore, we restricted our experiments to a time window between P21 - P25, where phenotypic abnormalities were still minimal and average body weights between the experimental cohorts indistinguishable. Consistent with a functional role of Kif1a in auditory perception, we found elevated ABR thresholds in homozygous *Kif1a^lgdg^* mutants compared to heterozygous and wild-type litter mates (Figure 6A). Moreover, post-hoc immunohistochemical analysis revealed normal synapse counts in the mid-apical cochlear turns of the mutants (Wt_mean_: 17.68 ± 1.15 per IHC; *Kif1a^lgdg^*_mean_: 17.20 ± 0.88, p=0.775; Figure 6B, C), although ribbon volumes of *Kif1a^lgdg^* mice were significantly reduced (Wt_median_ 0.063 μm^3^ IQR 0.044-0.086; *Kif1a^lgdg^*_median_ 0.053 μm^3^ IQR 0.035-0.081; p<0.0001; Figure 6D). This is indicative of defective synapse assembly or structural maintenance during maturation. To investigate this latter process in more detail, we expanded our analysis to the early stages of postnatal development (Figure 7): At P5, the number of synaptically-engaged ribbons in Kif1a^lgdg^ mice was indeed lower than of their Wt littermates, while no change could be observed in the cytosolic precursor fraction (synaptic Wt_mean_: 54.42 ± 0.78 per IHC; synaptic *Kif1a^lgdg^*_mean_: 49.88 ± 1.66, p=0.0085; cytosolic Wt_mean_: 0.71 ± 0.11; cytosolic *Kif1a^lgdg^*_mean_: 0.51 ± 0.10; p=0.368; Figure 7A,B). At this developmental age, we furthermore found that ribbon volume – synaptic as well as cytosolic – was already significantly reduced in *Kif1a^lgdg^*mice (synaptic Wt_median_ 0.036 μm^3^ IQR 0.014-0.063; synaptic *Kif1a^lgdg^*_median_ 0.029 μm^3^ IQR 0.011-0.050, p<0.0001; cytosolic Wt_median_ 0.012 μm^3^ IQR 0.003-0.030; cytosolic *Kif1a^lgdg^* _median_ 0.005 μm^3^ IQR 0.002-0.019; p=0.0065; Figure 7C), thereby suggesting an overall decline in ribbon precursor volume acquisition that is carried on towards adulthood. When assessing even younger P3 mice, we found comparable numbers of synaptically-engaged ribbons (synaptic Wt_mean_: 51.57 ± 1.72; synaptic *Kif1a^lgdg^*_mean_: 51.75 ± 1.63, p=0.2678), but a trend towards reduced cytosolic ribbon precursor counts (cytosolic Wt_mean_: 0.97 ± 0.084; cytosolic *Kif1a^lgdg^*_mean_: 0.69 ± 0.112; p=0.0617; Figure 7D,E). Moreover, at this slightly earlier stage of postnatal development, the difference in synaptic ribbon volume failed to reach statistical significance, but displayed a trend towards volume reduction (synaptic Wt_median_ 0.032 μm^3^ IQR 0.012-0.061; synaptic *Kif1a^lgdg^*_median_ 0.031 μm^3^ IQR 0.012-0.058, p=0.0747; Figure 7F). Remarkably, the cytosolic precursor volume of *Kif1a^lgdg^* mice was significantly smaller compared to littermates (cytosolic Wt_median_ 0.011 μm^3^ IQR 0.003-0.027; cytosolic *Kif1a^lgdg^* _median_ 0.005 μm^3^ IQR 0.002-0.016; p=0.0134), thereby indicating a potential primary defect in cytosolic RIBEYE accumulation.

**Figure 6:**
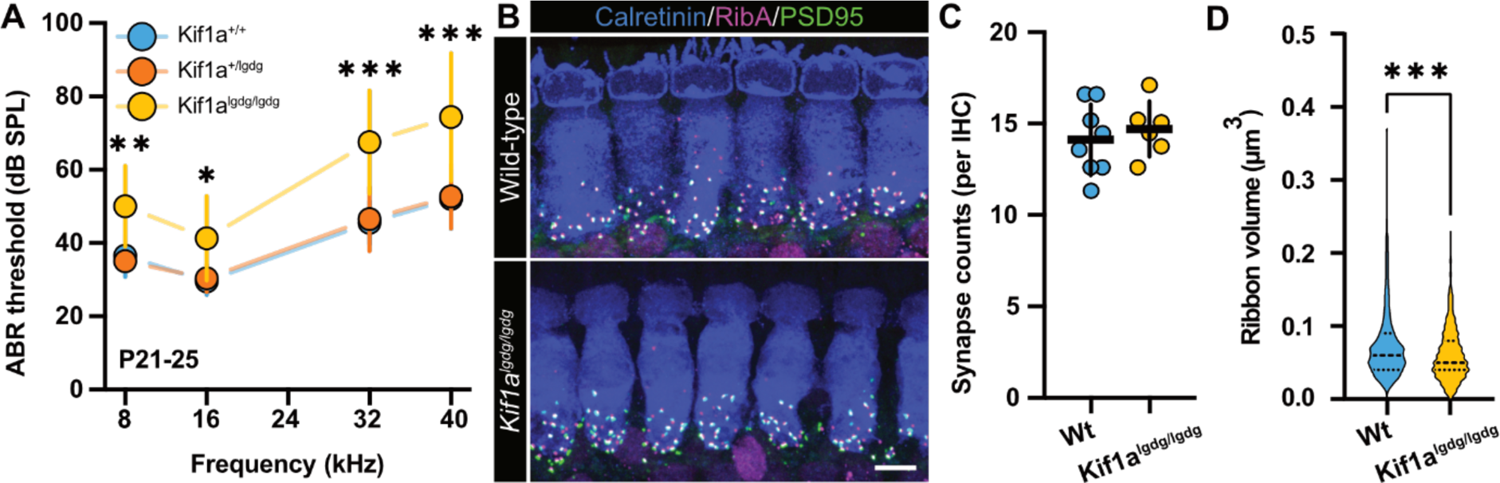
Kif1a is required for hearing and adequate IHC ribbon synapse volume acquisition. **A** Auditory brainstem responses (ABR) of P21-P25 mice carrying the *Kif1a^lgdg/lgdg^* mutation, compared to Wt and heterozygous littermates (*Kif1a^+/lgdg^*). Homozygous *Kif1a^lgdg/lgdg^* mutants displayed a moderate ∼10-20 dB increase in ABR thresholds for all tested frequencies, while the heterozygous mice showed intact hearing. n(Wt) = 10; n(Kif1a^+/lgdg^) =13; n(Kif1a^lgdg/lgdg^) = 8. Shown are means ± SD. **B** Representative confocal maximum projections of acutely-dissected organs of Corti of P21-P25 Wt littermates and *Kif1a^lgdg/lgdg^* mice, immunohistochemically labeled for RIBEYE (magenta), PSD95 (green) and IHC context (Calretinin, blue). **C** The number of ribbon synapses is comparable between mature *Kif1a^lgdg/lgdg^* mice and Wt littermates. **D** Ribbon volume of *Kif1a^lgdg/lgdg^* mice is reduced compared to Wt littermates.

**Figure 7:**
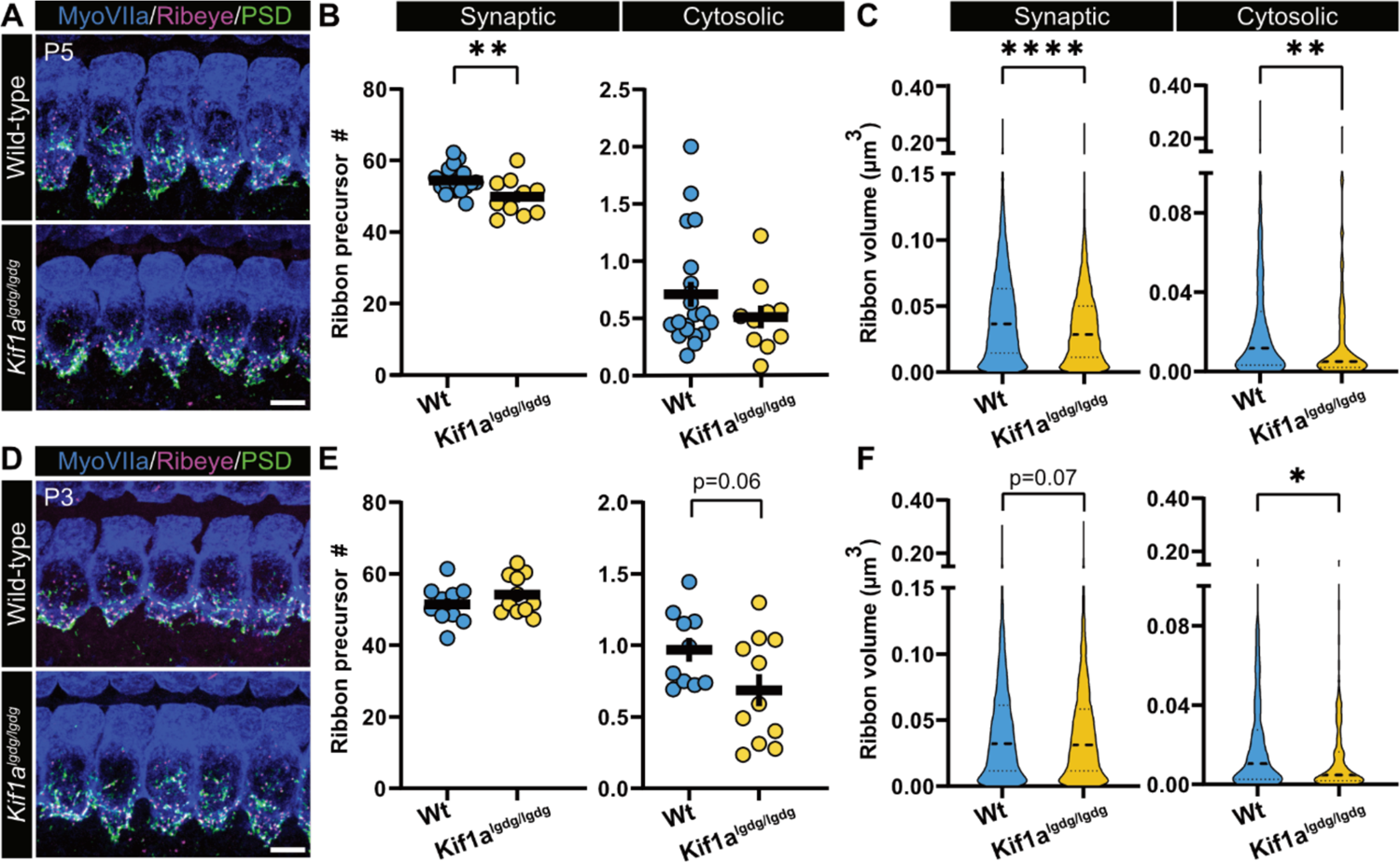
Impaired synaptic maturation in developing IHCs of *Kif1a^lgdg^* mutants. **A** Representative confocal maximum projections of acutely-dissected organs of Corti from P5 *Kif1a^lgdg/lgdg^* mice, immunohistochemically labeled for RIBEYE (magenta), PSD95(green) and IHC context (Myosin VIIa, blue). **B** The number of ribbon precursors that localize to the synapse is reduced in *Kif1a^lgdg/lgdg^* mice, whereas the number of cytosolic ribbons remains unaltered. **C** Ribbon volume in *Kif1a^lgdg/lgdg^* mice is reduced for synaptic as well as cytosolic ribbon precursors. **D** Representative confocal maximum projections of acutely-dissected organs of Corti from P3 *Kif1a^lgdg/lgdg^*mice, immunohistochemically labeled analogous to (A). **E** The number of synaptically-engaged and cytosolic ribbon precursors remains unaltered, although a trend towards reduction can be observed in the latter population. **F** Ribbon volumes in *Kif1a^lgdg/lgdg^*mice show a trend towards reduction for the synaptic population, while the cytosolic ribbon precursor fraction displays reduced volumes. Values represented either as individual datapoints with mean ± SEM, or as violin plots, with medians and the 25% and 75% interquartile range indicated with solid and dashed lines respectively. Statistical significance: Mann-Whitney U. * p<0.05, **p<0.01, ****p>0.0001. P5, N=15, n=30; P3, N=11, n=21. Scale bars: 5 μm.

## Discussion

The present work aimed to establish the molecular transport pathway of ribbon precursor spheres towards the developing presynaptic AZs of cochlear IHCs. For this purpose, AAV-transduced and genetically-as well as chemically-labelled IHCs were subjected to detailed live-cell and immunohistochemical analyses. Interrogation of cytoskeletal polarity revealed a highly polarized and strongly acetylated apico-basal MT network that enables longitudinal ribbon precursor trafficking to the presynaptic AZ and facilitates structural plasticity of ribbon precursors. Acute pharmacological disruption of the MT cytoskeleton impaired ribbon precursor velocity, displacement, directionality and volume acquisition – the latter via attenuation of the frequency of structural plasticity events between individual ribbon precursors and their functional interaction with MTs. In addition, phenotypic characterization of *Kif1a^lgdg^* mice revealed a moderate ABR phenotype and decreased IHC ribbon volumes that could already be detected during early postnatal development and thus implicates an essential role for kinesin-3 family member Kif1a in IHC synapse assembly and/or maturational refinement.

Therefore, together with our companion paper that analyzed ribbon precursor transport in zebrafish neuromast HCs, our combined data point towards an essential and evolutionary-conserved role of the polarized MT cytoskeleton and Kif1a-mediated transport in auditory ribbon synapse formation.

### Ribbon precursor translocation in IHCs is mediated by a MT-based transport system reminiscent of ‘slow’ axonal transport in neurons

To date, trafficking of synaptic components in small and compact cells – such as cochlear IHCs – remains poorly understood. In neurons, cargo trafficking has extensively been studied within the axon using radioisotopic pulse-labeling and live-cell microscopy experiments. Based on such work, targeted axonal transport was shown to employ a directional and multi-tiered trafficking system that comprises fast and slow delivery modes: while most SV proteins, neurotransmitters and transmembrane receptors are shuttled to their final destination via rapid MT- and molecular motor-based transport at rates ranging from ∼0.5-5 µm/s, non-membranous cytosolic proteins and soluble protein aggregates – including SV-associated proteins such as clathrin and synapsins, as well as cytoskeletal components – commonly travel at much lower velocities of ∼0.004-0.09 µm/s in a process that superficially resembles diffusion (Brown, 2000). However, rather than employing molecularly distinct mechanisms, the difference in speed of motion has been proposed to result from distinct frequencies and durations of the transient associations with the MT cytoskeleton, thus leading to saltatory ‘stop-and-go’ motility with alternating – in the latter case often prolonged – stationary periods and transient spurts of MT-based transport. On molecular level, this behavior appears to involve dynamic associations with components of the fast transport pathway, such as short-lived interactions with anterogradely trafficked SVs that produces an ‘anterogradely biased flow’ towards the synapse (Scott et al., 2011; Tang et al., 2013). According to this model – and given the observed mean ribbon precursor velocities of ∼0.006 µm/s – intracellular transport of ribbon precursors clearly falls into the ‘slow’ category and should be characterized by the occurrence of extended periods of (sub-)diffusive behavior with brief directional spurts of rapid displacement along MTs. Compatible with such a hypothesis, we found a significant fraction of ribbon precursors to be mobile along MT tracks and detected three main trajectory types: (i) saltatory, supra-linear MT-associated tracks, (ii) gradual/continuous, supra-linear MT-associated tracks, and (iii) non-directional, often spatially-confined tracks that likely represent membrane-anchored ribbons that reside at the presynaptic AZ. Interestingly, of the MT-associated tracks, the slow continuous mode presented the most prevalent category and – similar to the saltatory displacement mode – displayed velocity profiles indicative of interrupted motor-based transport. Future studies will have to assess if the underlying molecular mechanisms between these pathways share common features or are molecularly distinct.

### Acute MT disruption impacts synapse formation during early postnatal development

Consistent with a contribution of the MT network in ribbon precursor transport, acute pharmacological MT destabilization with nocodazole attenuated precursor velocity and displacement within IHCs. In particular, nocodazole treatment exerted prominent detrimental effects on precursor motion, as the fraction of supralinear trajectories was starkly reduced and trajectorial asymmetry of the low velocity precursors lost upon nocodazole application. Both findings are compatible with impaired directed transport. In contrast, fast-paced precursors, which likely represent a mixed population of non-directional, comparatively rapidly diffusive particles and fast-displacing precursors that undergo brief bouts of active transport, were found to be less affected by MT destabilization. Yet, the overall rather subtle reduction in precursor velocity, as well as the partly maintained fast targeted translocation upon pharmacological MT destabilization, is an indicator of incomplete disruption of the IHC MT network. In fact, the herein observed posttranslational acetylation of IHC α-tubulin is known to attenuate nocodazole-dependent MT depolymerization and strongly facilitates mechanical rigidity against strand breaks (Eshun-Wilson et al., 2019; Piperno et al., 1987; Portran et al., 2017; Xu et al., 2017). This likely prevents extended MT depolymerization in our experiments, and thus limits the destabilizing effect of nocodazole mainly to the MT +ends (Vasquez et al., 1997). As the displacement of the slow-moving particles is most strongly affected, this could indicate a preference for slow anterograde transport of precursors specifically taking place along the dynamic MT +ends. Future experiments will have to resolve this issue.

### Directionality and mode of transport for IHC ribbon precursor delivery to the AZ

Our data support an essential role of MT-based transport in ribbon synapse assembly; however, the molecular link between ribbon precursors and MTs remains elusive. To identify the involved molecular motors, it was essential to first establish MT polarity since each major molecular motor class has a preferred directionality: while kinesins predominantly travel to the MT +end, dyneins move towards the –ends. In the present study, we observed apical CAMSAP2 immunolocalization in the IHC neck. Hence, it can be assumed that the vast majority of centrosomal and non-centrosomal MTs are anchored at the IHC apex and that the MT +ends grow towards the basolateral compartment. In support of this hypothesis, our companion paper used live-cell single particle tracking of +end binding EB3-GFP in zebrafish neuromast HCs and found that the vast majority of EB3 trajectories (∼75%) project into the basolateral compartment, thus confirming the MT growth direction towards the synaptic region. This arrangement therefore supports an evolutionarily-conserved mechanism in which +end directed kinesin motors facilitate the anterograde delivery of ribbon precursors, other structural AZ components and SVs to the presynaptic AZ of developing IHCs. Mechanistically, this could most likely be achieved via a direct or indirect precursor/MT association, for example via RIBEYE, as has been described for other cytosolic and structural proteins – including synapsin, clathrin and neurofilaments. In fact, clathrin ‘transport packets’, which display slightly smaller outer diameters (∼125 nm) than ribbon precursors, have been shown to travel along neuronal axons at velocities of 0.006-0.5 µm/s in a MT-dependent manner (Ganguly et al., 2021), thus offering a mode of transport that is well compatible with our mobility data. Alternatively, ribbon-associated SVs may act as precursor/MT adaptors, yet – due to the energetic inefficiency of such a connection via a flexible protein linker (i.e., the filamentous SV tether) ‘dragging’ the precursor through a highly viscous environment – such a mechanism seems rather unlikely and warrants future studies for clarification.

### The anterograde motor Kif1a facilitates synapse assembly in IHCs

We previously proposed that ribbon precursor transport may involve the MT +end directed motor Kif1a (Michanski et al., 2019). This hypothesis was based on the established function of Kif1a in SV precursor transport to the presynaptic compartment (Okada et al., 1995) and our own observations of Kif1a colocalization with ribbon precursors as well as the close physical proximity between ribbon-associated SVs and MTs. Since our above findings are generally compatible with such a hypothesis, we now sought to investigate ribbon synapse morphology and ABR thresholds in the *Kif1a^lgdg^* mutants. These measurements revealed an early onset hearing impairment in *Kif1a^lgdg^*mice. In these animals, ribbon volumes were reduced compared to wild-type littermates – a phenomenon that could already be observed in a separate cohort of early postnatal *Kif1a^lgdg^* mice and is hence consistent with impaired volume accumulation during developmental maturation. Moreover, these data are compatible with our companion paper, in which genetic disruption of *kif1aa* in zebrafish lateral line HCs produced a similar – yet more striking – phenotype, as ribbon precursor areas were significantly reduced and on top, fewer synapses formed overall. Interestingly, the authors further found that this phenotype was due to attenuated precursor fusogenicity, rather than impaired overall transport rates. Our data may also support such a scenario: Since ribbon counts were only transiently affected and the difference in precursor volumes was more pronounced in the later stages of developmental maturation, Kif1a appears to play a key role in the gradual accumulation of ribbon material and concomitant structural maintenance rather than solely supporting initial synapse formation. Future live-cell studies in *Kif1a^lgdg^* will hence be required to clarify the exact role of Kif1a in IHC synaptogenesis.

Considering that IHCs are small and compact cells, the observed decrease in ribbon volume at such an early age is remarkable, given that the hind limb phenotype – which involves long-distance axonal transport – only manifests in the 3^rd^ to 4^th^ postnatal week. Moreover, the mutation likely does not confer a complete loss of Kif1a function. Therefore, it can be assumed that Kif1a – while presumably not being the only motor involved in anterograde ribbon precursor transport – plays an important role that cannot entirely be compensated by motor redundancy. Here, potential candidates may include the anterograde kinesin-2 motor Kif3a, which has been found to associate with RIBEYE in multiple ribbon-bearing sensory systems (Muresan et al., 1999; Spiwoks-Becker et al., 2008; Uthaiah and Hudspeth, 2010). Thus, although methodologically demanding, future work should aim to determine the exact time course of Kif1a involvement and identify other relevant anterograde as well as retrograde motors.

Finally, regarding the elevation of ABR thresholds, it should be highlighted that – based on the established role of Kif1a in SV transport and its wide neuronal expression pattern – it can be expected that the observed *Kif1a^lgdg^* hearing impairment phenotype reflects a cumulative effect on acoustic perception that most certainly also involves other components of the ascending auditory pathway. Hence, future studies should dissect the hearing phenotype of these mice in greater detail.

### Ribbon precursors regularly undergo MT-dependent structural plasticity events

A surprising finding of our study is the striking structural plasticity of ribbon precursors: lineage tracing analysis revealed the frequent occurrence of fusion and fission events. At this point, it remains unclear if ‘fusions’ involve the collision and intermixing of individual precursors or rather reflect a transient and reversible interaction, for example via tethering of the same SVs. Nevertheless, these findings contrast previous assumptions of a purely unidirectional pathway that leads to the accumulation of ribbon precursor material at the developing AZ (Michanski et al., 2019). Rather, these observations indicate that balanced and *bi-directional* precursor plasticity is a crucial component of ribbon synapse assembly and essentially requires an intact MT cytoskeleton: Upon nocodazole-dependent MT destabilization, the frequency of both types of plasticity events was significantly reduced and resulted in abnormal ribbon volume accumulation. In contrast, genetic disruption of *Kif1a* led to an early-onset reduction in ribbon size, thereby offering insights into the importance of *balanced* transport mechanisms: when anterograde motors are impaired but putative retrograde pathways left intact, ribbons fail to adequately accumulate material – likely due to an induced over-representation of ribbon fission events. Future live-cell analyses in *Kif1a^lgdg^* mice should test this hypothesis. Interestingly, our data are partly consistent with the findings of our companion paper, which reports nocodazole-sensitive and Kif1aa-dependent ribbon precursor fusions in zebrafish neuromast hair cells, thereby indicating a evolutionarily-conserved MT-based mechanism underlying ribbon synapse formation.

To this end, the exact role of precursor fissions for IHC ribbon synapse development – together with its underlying molecular mechanism – remains to be determined, but given the seemingly strict apicobasal MT polarity likely involves MT –end directed retrograde motors of the dynein family. Future studies will be required to test this idea experimentally.

### Cytoskeletal roles in IHC synapse maturation

Our observations of ribbon precursor dynamics at the beginning of the second postnatal week suggest a redistribution of ribbon precursor material rather than clear long-distance apico-basal precursor translocation. Therefore, MT-based transport likely contributes to the structural refinement process of the maturing AZ. Here, it is tempting to speculate that upon maturational pruning of individual synaptic contacts, detached ribbons are locally trafficked to adjacent AZs rather than being proteasomally degraded. While this hypothesis will have to be experimentally validated, it is compatible with previous electron microscopy studies that showed floating ribbon precursors in the IHC cytoplasm not only around the time of initial AZ assembly, but also towards hearing onset, when the initial establishment of synaptic contact sites should have fully concluded (Michanski et al., 2019; Sobkowicz et al., 1986, 1982; Wong et al., 2014). Such local re-distribution of surplus ribbons might additionally be supported by the cortical F-actin cytoskeleton, which has been suggested to form ‘cage-like’ structures at IHC AZs, possibly constituting diffusion barriers that control Ca^2+^-dependent SV exocytosis (Guillet et al., 2016; Vincent et al., 2015). In addition to this direct role in SV release, it is also conceivable that an actin/myosin-based transport system plays a complementary role to MT/Kif1a-based mechanisms in the local re-distribution of detached ribbon precursors and hence, would present an interesting topic for future studies.

In summary, our data shed light on the still poorly understood mechanisms underlying auditory ribbon synapse formation. In recent years, various studies have shown that upon hair cell loss – e.g., through traumatic noise, ototoxicity or degeneration – supporting cells can be reprogrammed into HCs. Yet, such ectopic HCs need to also be adequately innervated. Therefore, a fundamental understanding of IHC ribbon synaptogenesis is an essential prerequisite for the design of future restorative therapies to regenerate lost auditory synapses.

## Materials & Methods

### Animals

The recessive *Kif1a* leg dragger (*Kif1a^lgdg^*) mutation (RRID: IMSR_JAX:016894) was isolated at The Jackson Laboratory and mapped as a C to T point transition at position 93,076,218 bp (GRCm38/mm10), causing a L181F amino acid change (MGI Direct Data Submission J:229662). For this work, this strain was in a mixed genetic background after breeding with C57BL/6J animals to improve health and lifespan. All animal experiments were conducted according to national, regional and institutional guidelines of either Göttingen, Lower Saxony, Germany for wild type C57Bl6/J mice (WT) and Ai32-Vglut3-Cre knock-in mice (Ai32-VC-KI; (Chakrabarti et al., 2022)), or Bar Harbor, Maine, USA, for *Kif1a^lgdg^*mice. All experiments were approved by the respective animal welfare officers. Mice of either sex between age postnatal day (P)3 and 25 were sacrificed by decapitation for either acute dissection and fixation of the organ of Corti, or preparation of organotypic explant cultures for live-cell imaging. Mice past P5 were euthanized via cervical dislocation at The Jackson Laboratory.

### Preparation of mouse organotypic cultures of the organ of Corti

Organotypic cultures of the organ of Corti were prepared from neonatal WT and Ai32-VC-KI mice (P5-P7). Preparation procedures were based on (Vogl et al., 2015), with adaptation of the culturing medium to Neurobasal Medium (#12349-0.15, Gibco) supplemented with GlutaMAX (1%, #35050-061, Gibco), B27 Plus Supplement (2%, #A35828-01, Gibco), and Ampicillin (1.5 μg/mL). In brief, the apical-medial turn of the organ of Corti was dissected from the mouse cochlea and mounted on either 1.5 thickness high-precision coverslips or glass bottom Petri dish inserts (P35G-1.5-14-C, MatTek), coated with CellTak (#354240, Corning, 1:8 solution in NaHCO3). Subsequently, organotypic cultures were submerged in 2 ml culturing medium in a 35 mm Petri dish and incubated at 37°C, 5% CO_2_ for up to two days *in vitro* (DIV2).

### Molecular cloning of the construct, virus production and purification

Transgene expression of the RIBEYE (NCBI Reference Sequence: NC_000073.7) with EGFP as fusion protein was promoted by the hybrid promotor hCMV/HBA (human cytomegalovirus immediate early enhancer, human beta-actin promotor). The Woodchuck Hepatitis Virus Posttranslational Regulatory Element (WPRE) and the bovine growth hormone (bGH) polyadenylation sequence were included in the construct (pAAV) to enhance transcription and improve the stability of the transcript. The same promoter was used for the tdTomato version of the RIBEYE (RIBEYE-tdTomato) generated by molecular cloning performed by AgeI and Sal I enzymatic digestion followed by a ligation procedure. Both constructs were validated by sequencing using the Sanger DNA sequencing methodology. The generated RIBEYE-GFP and RIBEYE-tdTomato constructs were packaged into AAV9-PHP.B and AAV9-PHP.eB, respectively (Chan et al., 2017). PHP.(e)B particles were generated using our standard AAV purification procedure previously described in more detail in (Huet and Rankovic, 2021). In brief, triple transfection of HEK-293T cells was performed using pHelper plasmid (TaKaRa/Clontech), trans-plasmid providing viral capsid PHP.(e)B (a generous gift from Viviana Gradinaru (Addgene plasmid #103005) and cis plasmid providing hCMV/HBA_wtRIBEYE-EGFP or tdTomato. PHP.(e)B viral particles were harvested 72 h after transfection from the medium and 120 h after transfection from cells and the medium. Precipitation of the viral particles from the medium was done with 40% polyethylene glycol 8000 (Acros Organics, Germany) in 500 mM NaCl for 2 h at 4°C. Both, precipitate and cells were lysed in high salt buffer (500mM NaCl, 2mM MgCl 2, 40mM Tris-HCl pH 8,0) and non-viral DNA was degraded using salt-activated nuclease (SAN, Arcticzymes, USA). Afterward, the cell lysates were clarified by centrifugation at 2,000 g for 10 min and then purified over iodixanol (Optiprep, Axis Shield, Norway) step gradients (15, 25, 40, and 60%) at 350000xg for 2.25 h (Grieger et al., 2006; Zolotukhin et al., 1999). Finally, viral particles were concentrated using Amicon filters (EMD, UFC910024) and formulated in sterile phosphate-buffered saline (PBS) supplemented with 0.001% Pluronic F-68 (Gibco, Germany). The virus titer of RIBEYE-EGFP was 4.70*10^12^ - 5.37*10^12^ genome copies/ml and of RIBEYE-tdTomato was 2.14*10^12^ genome copies/ml, determined according to the manufacturer’s instructions by determining the number of DNase I resistant vg using qPCR (StepOne, Applied Biosystems) and AAV titration kit (TaKaRa/Clontech). The purity of produced viruses was routinely checked by silver staining (Pierce, Germany) after gel electrophoresis (NovexTM 4–12% Tris-Glycine, Thermo Fisher Scientific) according to the manufacturer’s instructions viral stocks were kept at −80°C until the injection day.

#### *In vivo* AAV injections

Mice were injected P4-6 using the round window approach as described in earlier studies (Huet and Rankovic, 2021; Rankovic et al., 2021). In brief, anesthesia was established with isoflurane (5% for induction, 2–3% for maintenance, frequent testing of the absence of hind-limb withdrawal reflex). For analgesia, buprenorphine (0.1 mg/kg body weight, injection 30 minutes before surgery) and carprofen (5 mg/kg body weight, applied during and 1-day post-surgery) were applied subcutaneous and xylocain (10 mg spray) locally. Body temperature was maintained by placing the animal on a remote-controlled custom-built heating blanket. Following a retro-auricular approach, the facial nerve was exposed in order to determine where to puncture the cartilaginous bulla with the injection pipette and target the scala tympani where virus suspension (∼2 μl, corresponding to 9.4*10^9^ – 1.074*10^10^ AAV particles (RIBEYE-EGFP) and 4.28*10^9^ AAV particles (RIBEYE-tdTomato)) was injected. Following the injection, the endogenous tissue was relocated, and the surgical situs was closed by suturing the skin. One day after injections, mice were used for organ of Corti organotypic culture preparations and subsequent live-cell imaging and immunohistochemistry.

### Live-cell labeling of the IHC MT cytoskeleton

To fluorescently label the IHC MT cytoskeleton, a small region of the organotypic culture (approximately 10 IHCs) was mechanically cleared of outer hair cells (OHC) and supporting cells using the established glass micropipette-based cleaning technique for electrophysiology experiments on auditory IHCs – e.g., in (Vogl et al., 2015). This includes the use of various glass pipettes with decreasing μm-sized tip diameters to aspirate surrounding cell types by gentle suction – including the OHCs, outer as well as inner pillar cells and phalangeal cells. Thereafter, the MT labeling dye SPY-555-tubulin (#SC203, Spirochrome) was applied to the cleaned culture (1 μM), and treated cultures were then incubated for 6 hours at 37°C, 5% CO2. This dye application method facilitated optimal tissue penetration and thereby IHC targeted MT labeling with minimal optical interference from strongly tubulin-expressing adjacent supporting cells (especially inner pillar cells).

In preparation for live-cell imaging, coverslip-attached organotypic cultures were inverted and placed inside the insert of a glass bottom culturing dish (P35G-1.5-14-C, MatTek), thereby creating a thin slice-like section between the two glass coverslips. This allowed for direct, top-down accessibility of the IHCs and enabled visualization of the fine MT network by minimizing the objective-to-tissue working distance and circumventing any tissue-induced aberrations arising from acquisition through the dense basilar membrane.

### Manipulation of the IHC MT cytoskeleton in vitro

Nocodazole (Nocodazole Ready Made Solution, #SML1665, Sigma-Aldrich) was applied to P6 or P7 Ai32-VC-KI organotypic cultures of the organ of Corti on DIV2, for 3 hours at a final concentration of either 1 μM nocodazole or vehicle (DMSO). Thereafter, treated cultures were used for live-cell timelapse imaging experiments.

### Fixation and immunohistochemistry of organotypically-cultured or acutely-dissected organs of Corti

Immunohistochemistry to assess the volume and synaptic engagement of ribbon precursors included organotypic cultures of WT and Ai32-VC-KI mice, as well as acute dissections of *Kif1a^lgdg^* mice and wild-type (Wt) littermates. The organotypic cultures were fixed in 4% formaldehyde for 15 min on ice. Acutely dissected cochleae of *Kif1a^lgdg^* and Wt littermates were fixed in 4% formaldehyde for one hour on ice, stored in PBS/0.02% sodium azide and shipped to Germany on ice for dissection and further processing for immunohistochemistry.

Fixed explant cultures and acutely dissected cochleae were permeabilized in PBS + 0.5% Triton-X100 for 30 min, and thereafter incubated in blocking solution (PBS + 0.5% Triton-X100 + 10% normal goat serum) for one hour. Incubation in primary, as well as secondary or directly-conjugated antibodies was performed in blocking solution, for two hours at room temperature protected from light. Samples on coverslips were mounted on glass slides, whereas samples in glass bottom dishes were covered with a coverslip using ProLong Gold Antifade reagent (#P36984, Invitrogen).

Immunohistochemistry experiments to label the IHC cytoskeleton included organotypic cultures and acute dissections of WT mice. The organs of Corti were extracted for 3 minutes using prewarmed extraction buffer (Jansen et al., 2023), and subsequently fixed with 4% formaldehyde for 30 minutes, at 37°C. Permeabilization, blocking and antibody incubation was done as described above. The following antibodies were used in this study: anti-Calretinin (Chicken, SySy, #214106), anti-RIBEYE-A (Rabbit, SySy, #192103); anti-acetylated tubulin (Mouse IgG2b, Sigma, #T7451); anti-CAMSAP2 (Rabbit, Proteintech, #17880-1-AP); anti-Myosin VIIa (Mouse IgG1, Developmental Studies Hybridoma Bank, #MYO7A 138-1); and anti-β-3-tubulin (Tuj-1; Mouse IgG2a, BioLegend, #801202) and a fluorescently-conjugated nanobody directed against PSD-95 (Fluo-Tag-X2, Alexa Fluor647 conjugated, NanoTag, #N3702-AF647). For final visualization standard AlexaFluor - 488, - 594 and - 647-conjugated secondary antibodies were used (ThermoFisher Scientific).

### Image acquisition

For optimal optical resolution of MTs, live-cell imaging experiments for ribbon precursors in MT context were performed on an Abberior Instruments Expert Line STED microscope – operated in confocal laser scanning mode and equipped with a 60x/NA 1.20 water immersion objective. Environmental control (37°C, 5% CO_2_) was achieved with a top mount on-stage incubator (Okolab uno stage-top incubator, H391-Olympus-IX-SUSP 2015). Regions of interest were selected for low to moderate RIBEYE-GFP expression, strong SPY555-tubulin labeling intensity, IHC orientation and healthy IHC morphology. Timelapse images were acquired over a period of 30-75 minutes of continuous imaging at maximum acquisition speed. Depending on IHC orientation, the required axial depth of the imaging stack varied; hence, the image acquisition intervals varied between 35 and 90 seconds per stack.

Live-cell imaging experiments for ribbon precursors in IHC cellular context were conducted at a Nikon Eclipse Ti Andor Spinning Disk confocal imaging setup, 60x water immersion, NA 1.20, under environmentally controlled conditions (37°C, 5% CO_2_, Okolab Bold Line Cage Incubator). Regions of interest were selected for low to moderate RIBEYE-tdTomato expression and healthy IHC morphology. Timelapse images were acquired for 40 minutes, by continuous z-stack acquisition at intervals of exactly 50 seconds.

Immunolabelled samples were imaged using an Abberior Instruments Expert Line STED microscope. Z-stacks were acquired in confocal mode using a 100x/NA 1.4 oil immersion objective.

### Image processing and analysis

Timelapse images were corrected for photobleaching (BleachCorrection (Miura, 2020), FIJI/ ImageJ, 2.3.0/1.53q), and physical drift (IMARIS; Oxford instruments, 9.6.1). Ribbon precursor particles were detected and traced using the Spots particle tracking function, under the lineage tracing algorithm (0.5 seed point diameter, 1.0 PSF correction, background subtraction, 20 seed point quality threshold, 45 region border growing, lineage tracing, 1.5 μm maximum distance, 0 maximum time gap). Volume assessment of ribbon precursors from timelapse images was performed using the IMARIS Surface rendering function (0.1 surface detail, local background subtraction, 0.28 μm largest sphere diameter, 0.3 split surface seed points, 10 quality filter, filter closest distance to Spots=Ribbons).

Confocal images of fixed tissue were analyzed for ribbon number and volume using the IMARIS Surface rendering function. Ribbon precursors (0.054 surface detail, local background subtraction, 0.28 μm largest sphere diameter, 0.15 split surfaces, 3.0 seed point diameter, 3.0 quality filter) were classified based on their proximity to the 3D rendering of the PSD (0.08-0.16 surface detail, local background subtraction, 0.6 μm largest sphere diameter, 3.0 quality filter), and classified as either synaptically-engaged (within 500 nm surface-to-surface distance to the PSD), or cytosolic ribbons. Normalization of ribbon volumes and numbers was performed by dividing of the ribbon precursor values of pharmacologically treated conditions by the mean of the respective control.

For the assessment of three-dimensional ribbon precursor displacement, positions of precursors were determined by their center of mass, using the IMARIS Spots function. The mean squared displacement (MSD) was then calculated based on the extracted xyzt coordinates for individual precursor trajectories with the transversed displacement averaged over progressive imaging frames / time steps (τ). In the extracted MSD, N is the number of data points in a trajectory, Δτ is the time interval per imaging frame, and x, y and z are the ribbon precursor coordinates.

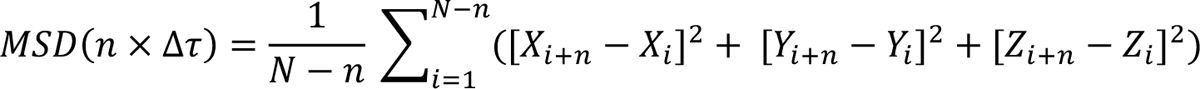

The MSD curve was then fitted using a least squares fit to determine the exponent α, as well as parameter K.

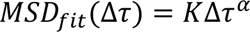

The asymmetry measure, originally introduced by (Huet et al., 2006) quantifies the anisotropy of the particle motion. It is calculated from the eigenvalues of the gyration tensor, which is the dimensional counterpart of the MSD. For symmetric motion, the eigenvalues of the gyration tensor will grow symmetric for higher time intervals. However, for asymmetric motion, these eigenvalues become unequal. With R_1_, R_2_ and R_3_ being the square roots of the eigenvalues of the gyration tensor, also known as the gyration radii, the asymmetry measure is:

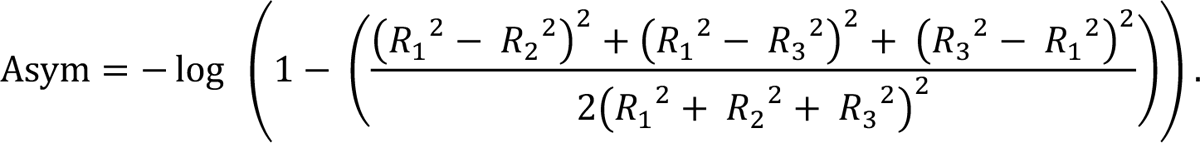

To conduct the asymmetry analysis, tracks generated by the lineage tracing algorithm were split at timepoints of plasticity events; then filtered for the ‘main trajectories’ to only included tracks spanning more than 10 imaging frames (>500 s), and for mobility, excluding stationary tracks of velocities below 0.001 μm/s.

The variability measure shown as shaded areas for the calculated MSD and asymmetry were determined to encapsulate the corresponding values from the symmetric 95% of the bootstrap samples. For all Δτ, 100 bootstrap samples were sampled with replacement from all spatial steps independently (Efron and Hastie, 2021).

The MSD and asymmetry measure were determined with original scripts in Python. The asymmetry measure was adapted for three-dimensional space.

### Statistics

Statistical analysis was performed in Prism8 (GraphPad, San Diego, CA). To assess the normality of the distributions, a D’Agostino-Pearson’s test was used. Statistical significance between two groups was then determined with an unpaired Student’s *t-test* or Mann-Whitney U test; for comparison between multiple groups a Kruskal-Wallis test was performed in combination with a multiple comparisons Dunn’s post-hoc. Values in the text are presented as classification_median_, classification_IQR_ (inter quartile range), or as mean ± SEM or SD as stated in the respective text section.

### Auditory Brainstem Response (ABR) recordings

All tests were performed in a sound-attenuating chamber, and body temperature of the anesthetized animals was maintained at 37°C using a heating pad (FHC Inc.). Animals of both sexes between P21 and P25 were anesthetized using a mix of ketamine and xylazine (1 mg and 0.8 mg per 10 g of body weight, respectively) and tested using the RZ6 Multi-I/O Processor System coupled to the RA4PA 4-channel Medusa Amplifier (Tucker-Davis Technologies). ABRs were recorded after binaural stimulation in an open field by tone bursts at 8, 16, 32, and 40 kHz generated at 21 stimuli/second. A waveform for each frequency/dB level was produced by averaging the responses from 512 stimuli. Subdermal needles were used as electrodes, with the active electrode inserted at the cranial vertex, the reference electrode under the left ear and the ground electrode at the right thigh. ABR thresholds were obtained for each frequency by reducing the sound pressure level (SPL) by 5 decibels (dB) between 90 and 20 dB to identify the lowest level at which an ABR waveform could be recognized. We compared waveforms by simultaneously displaying 3 or more dB levels on screen at the same time.

## Acknowledgements

We would like to thank Christiane Senger-Freitag and Sandra Gerke for expert technical support and Katie Kindt for the helpful discussions and comments on the manuscript. Moreover, we would like to express our gratitude to Cathleen Lutz (The Jackson Laboratory) for sharing the *Kif1a^lgdg^* strain in a mixed genetic background to improve animal health and Tina Pangrsič for providing Ai32-VC-KI mice. This work was funded by project B08 of the Collaborative Research Center 889 ‘*Cellular Mechanisms of Sensory Processing*’ of the German Research Foundation (to CV) and an *Otto Creutzfeldt Fellowship* of the Elisabeth and Helmut Uhl Foundation (to CV). BT was supported by grants R01 DC015242 and DC018304 from the National Institute on Deafness and Other Communication Disorders (NIDCD).

## Author contributions

RAV and CV designed the experiments, RAV performed live-cell imaging, immunohistochemistry and data analysis. CV performed immunohistochemistry and data analysis. AJ and BT maintained the *Kif1a* mouse colony, collected tissue for immunohistochemistry and performed ABR analysis. MS and FW analyzed data. VR designed and generated AAVs and performed intra-cochlear injections. RAV and CV wrote the paper and generated the Figures. All co-authors revised the manuscript.

## Supplemental Figures

**Supplemental Figure 2-S1:**
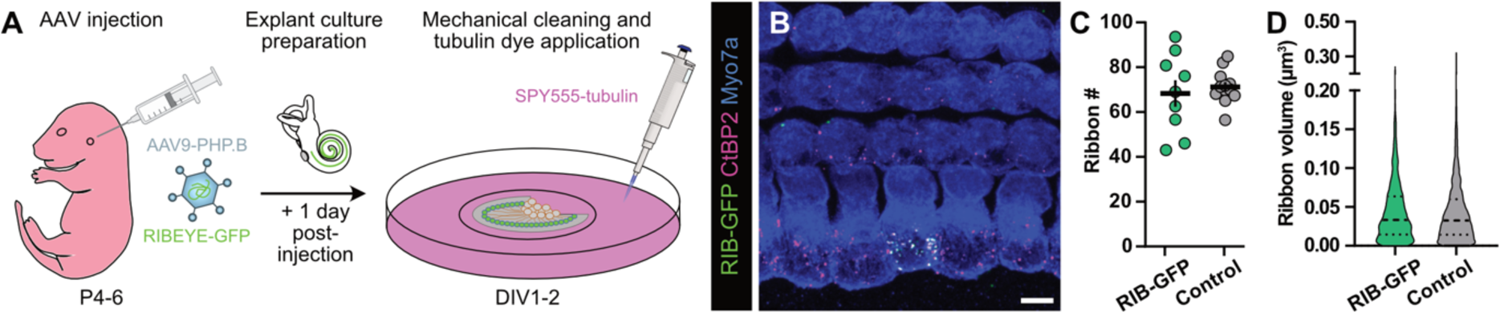
Experimental paradigm and effects of short-term RIBEYE-GFP overexpression on ribbon count and volumes. **A** Wild-type mouse pups were injected with an AAV encoding RIBEYE-GFP at postnatal day P4-6. One day after transduction, organ of Corti explant cultures were prepared and – after additional one to two days *in vitro* (DIV) – mechanically-cleaned and incubated with the MT dye SPY555-tubulin. **B** Representative maximum projection of a confocal z-stack showing a transduced IHC, which expresses RIBEYE-GFP (green). Please note the colocalization with the ribbon marker CtBP2 (magenta). **C-D** Both, ribbon counts (C) and volumes (D) were indistinguishable between RIBEYE-GFP transduced and neighboring non-transduced IHCs, suggesting appropriate integration of the fluorescent construct into endogenous ribbons while not displaying any obvious overexpression artifacts. No statistical significances detected (Mann-Whitney U test). RIBEYE-GFP transduced: N=9, n=9; Control non-transduced: N=12, n=14. Scale bar: 5 μm.

**Supplemental Figure 5-S1:**
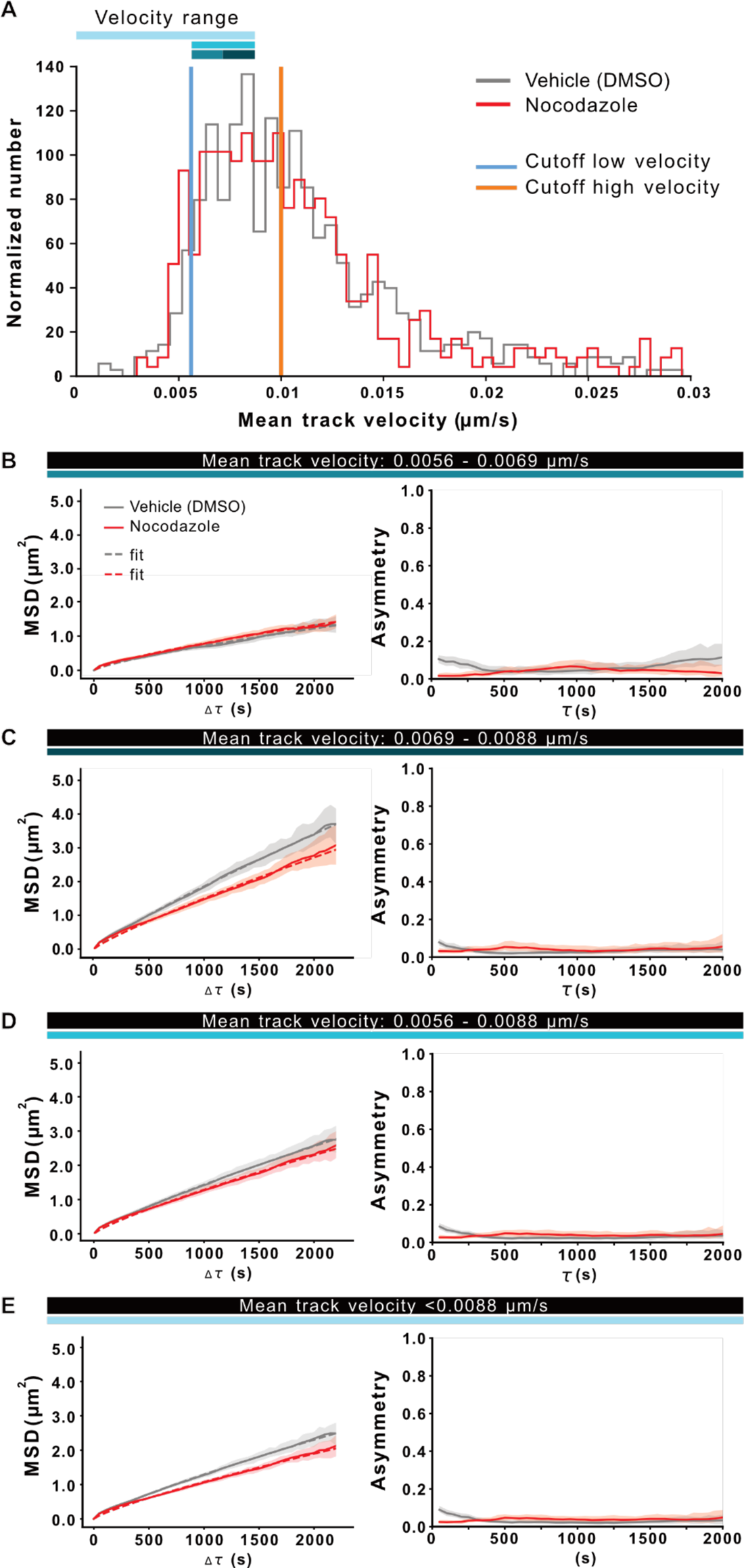
Dilution of nocodazole effects in faster-displacing ribbon precursor populations. **A** Reproduction of the same dataset as in Figure 5C: Shown is the distribution of the mean track velocity for precursors in vehicle- and nocodazole-treated IHCs. Indicated are the used cutoffs to selectively analyze trajectories with a low (blue) and high velocity (orange) displacement. Color-coded bars indicate the different velocity ranges displayed in B-E. **B-E** MSD analysis (left panels) and asymmetry assessment (right panels) of trajectories with a mean velocity within the range 0.0056 – 0.0069 μm/s (B), 0.0069 – 0.0088 μm/s (C), 0.0056 – 0.0088 μm/s (D), and below 0.0088 μm/s (E). The inclusion of a moderate-to-fast displacing population of ribbon precursor trajectories dilutes the reducing effect of nocodazole on ribbon precursor displacement and directionality present in the low velocity trajectories (below 0.0056 μm/s, as seen in Figure 5D). **B-D** Trajectories with a velocity between 0.0056 and 0.0088 μm/s do not appear to be subjected to directed transport in control, nor nocodazole-treated conditions. **E** Addition of the 0.0056 – 0.0088 μm/s velocity range to the low velocity cutoff range (<0.0056 μm/s) largely eliminates the distinction in 3D displacement and directed transport of ribbon precursors between nocodazole-treated and control IHCs.

